# High current optogenetic channels for stimulation and inhibition of primary rat cortical neurons

**DOI:** 10.1101/145441

**Authors:** Lei Jin, Eike Frank Joest, Wenfang Li, Shiqiang Gao, Andreas Offenhäusser, Vanessa Maybeck

## Abstract

ChR2-XXL and GtACR1 are currently the cation and anion ends of the optogenetic single channel current range. These were used in primary rat cortical neurons *in vitro* to manipulate neuronal firing patterns. ChR2-XXL provides high cation currents via elevated light sensitivity and a prolonged open state. Stimulating ChR2-XXL expressing putative presynaptic neurons induced neurotransmission. Moreover, stable depolarisation block could be generated in single neurons using ChR2-XXL, proving that ChR2-XXL is a promising candidate for *in vivo* applications of optogenetics, for example to treat peripheral neuropathic pain. We also addressed an anion channelrhodopsin (GtACR1) for the next generation of optogenetic neuronal inhibition in primary rat cortical neurons. GtACR1‘s light-gated chloride conduction was verified in primary neurons and the efficient photoinhibition of action potentials, including spontaneous activity, was shown. Our data also implies that the chloride concentration in neurons decreases during neural development. In both cases, we find surprising applications of these high current channels. For ChR2-XXL inhibition and stimulation are possible, while for GtACR1 the role of Cl^−^during neural development becomes a new optogenetic target.

## Introduction

The past discovery of Channelrhodopsin (ChR) 1/2 ^1,2^ enabled a broad range of novel neurophysiological experiments using optogenetics ^3,4^. The corresponding optogenetic applications are commonly limited by the functional expression in a particular model system or the rhodopsin’s features, like distinct specificity for particular ions. To reach a high level of expression and the corresponding number of functionally produced rhodopsins, the supplementation of the chromophore all-trans-vitamin-a-aldehyde (retinal) has proved to be helpful ^5^ and under some conditions the supplementation is even obligatory ^3^. Recently a ChR2 mutant called ChR2-XXL was published to overcome this limitation of optogenetics with strikingly high expression and slow closing kinetics enabling high photocurrents even under low light and retinal conditions ^6^. Similar to its precursor, ChR2-XXL is a relatively unselective proton and cation channel enabling neuronal firing during illumination with blue light (**Fig. 1a**). The past optogenetic approaches to inhibit neuronal activity have been based on light-driven pumps and are thus less efficient. Recent studies describe the discovery of Anion Channelrhodopsins (ACRs) ^7^ as an alternative. Within this class, GtACR1 seems to have a relatively strict anion conductance and offers the possibility of efficient light gated neuronal silencing (**Fig. 1a**). ChR2-XXL already proved to be a powerful optogenetic tool in *Drosophila Melanogaster*. Here we further tested the two high current optogenetic channels (ChR2-XXL and GtACR1) in a mammalian system (primary rat cortical neurons) to propose what protocols are needed to control neural activity without any side effects on the cells. Both high ion current light-driven tools offer a next generation of neurophysiological optogenetics. Here we highlight consequent applications and set the field for advanced optogenetics.

**Figure 1.**
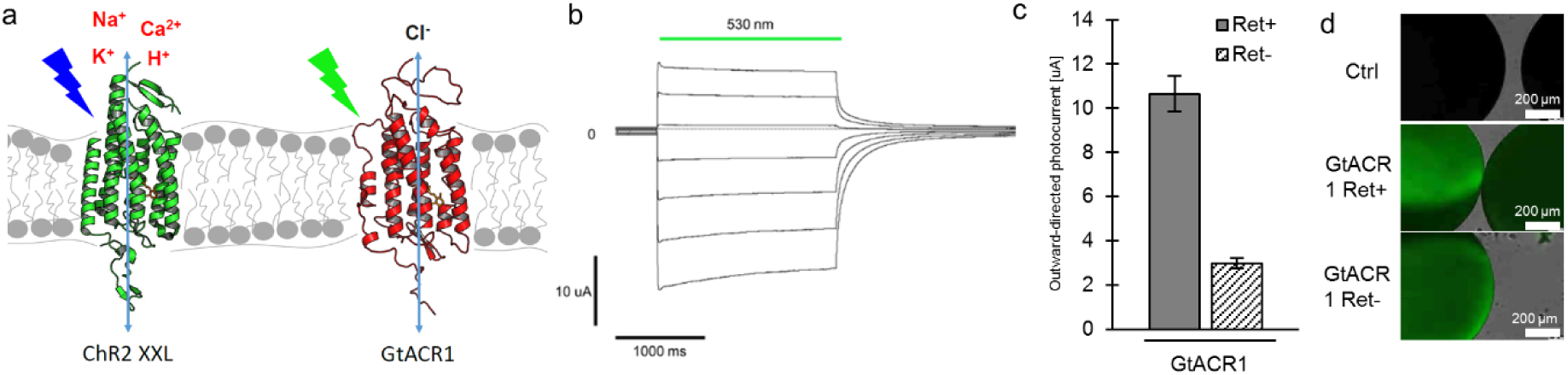
GtACR1 in *Xenopus laevis* oocytes. A) Schemes displaying the basic functional properties of ChR2-XXL (green) and GtACR1 (red). ChR2-XXL relatively unselectively conducts protons and cations, such as sodium, after irradiation with blue light (473 nm), whereas GtACR1 conducts anions, such as chloride, after irradiation with green light (530 nm) under physiological conditions. b) Photocurrents generated by a selected GtACR1 expressing *Xenopus laevis* (X.l.) oocyte 16 hours after injection of 5 ng cRNA (16 hpi). A saturating 530 nm light pulse (2 s) induces current as membrane potential is changed in +20 mV steps from -100 mV to +20 mV. c) Mean stationary photocurrent at -100 mV of oocytes (X.l.) expressing GtACR1 (16 hpi) after incubation in ND96 solution supplemented with 1 μM all-trans retinal (Ret+) or without (Ret-) (Ret+ *n*=10, Ret- *n*=9). d) Observed fluorescence of selected oocytes (X.l.) expressing the GtACR1::YFP construct 64 hpi and after incubation in Ret+ or Ret- ND96 (Ret+, Ret-). Electrophysiological recordings were done in ORI BaCl_2_ solutions pH 7.6. Rhodopsins were activated using a laser with corresponding wavelength at saturating intensities (GtACR1 530 nm = 18.2 mW/mm^2^). Each statistical evaluation is showing an individual paired experiment. Error bars indicate s.d.

Peripheral neuropathic pain originates in the peripheral nerve fibres, most commonly in the arms and legs ^8,9^. Scientists of neural engineering are seeking improved approaches that can be used to block pain fibres selectively and reversibly to alleviate episodes of debilitating chronic pain. An optogenetic approach ^2,10,11^ offers elegant ways to achieve this aim. Especially, ChR2-XXL ^6^, which can give rise to the largest photocurrents of all published ChRs and mutants with an increased light sensitivity more than 10,000-fold over wild-type ChR2 in *Drosophila* larvae. This results in gating by diffuse, ambient light and enables an efficient threshold surpassing depolarisation for non-invasive neuronal control. Moreover it is likely that a stable depolarisation block can be induced by photostimulating ChR2-XXL expressing neurons due to its large photocurrent ^12^, inactivating the voltage-gated sodium channels ^13^, or activation of calcium-activated K^+^ channels, resulting in a block of neuronal over-excitation ^14^.

Optogenetic studies of neurotransmission will play an important role in the fields of behaviour as well as neuronal disorders. Moreover, ChR2-XXL with its characteristic long open-state ^6^ likely allows future investigations of synaptic plasticity *in vivo*. Therefore, investigating neurotransmission by stimulating ChR2-XXL expressing cultured neurons will build a fundamental basis.

Channelrhodopsin-2 and variants thereof ^2^ were generated and characterised to become a toolbox for activating neural activity in response to light and are widely applied in neuroscience^3,4,15–17^. In contrast, inhibitory tools are less frequent in the literature and unequally effective for inhibiting neurons. However, recently discovered naturally occurring anion channelrhodopsins (ACRs) of the alga *Guillardia theta* might overcome these limitations and imbalance ^7^ by exclusive anion conductance. ACRs seem to be able to silence neuronal activity with high sensitivity and efficiency, ^7^ presumably due to their strict conduction of anions such as Cl^−^. Complete exclusion of protons and cations, results in their effective hyperpolarisation of the neural membrane, and usefulness in decoding brain functions.

Here, we present possible applications of high ion current optogenetics by expressing ChR2-XXL or GtACR1 in primary rat cortical neurons. We introduced optogenetic investigations of neurotransmission with ChR2-XXL in primary rat cortical neurons and moreover we investigated whether a stable depolarisation block can be successfully induced in ChR2-XXL expressing neurons. Furthermore, the function of GtACR1 was validated in primary cortical neurons by successfully achieving photoinhibition of spiking induced by a pulsed current as well as inhibition of spontaneous activity. Our findings will strongly facilitate and broaden these tools to investigate how the inhibitory or excitatory neurons control the behaviour. Aside from the efficiency of ChR2-XXL and GtACR1 in standard applications, these additional uses show that there are more areas of optogenetics to be developed using high current channels. These two high current optogenetic channels (ChR2-XXL and GtACR1) will provide a sufficient toolbox for optical control to decode neural circuits and understand basic brain function ^18,19^.

## Results

### GtACR1 or ChR2-XXL in *Xenopus laevis* oocytes

We validated GtACR1 by generating a GtACR1::YFP construct and injecting the cRNA into *Xenopus laevis* (*X.l*.) oocytes for TEVC recordings. High photocurrents were already detected 16 hours post injection (hpi) of 5 ng/oocyte cRNA with saturating 530 nm light pulses (**Fig. 1b**). Surprisingly, no obvious YFP signal was observed by fluorescence microscopy at this time, indicating a low GtACR1 expression level and relatively high single channel conductance. ChR2-XXL’s increased photocurrent is associated with a stronger expression level ^6,20^. To further validate these observations, photocurrents and the fluorescence of oocytes expressing GtACR1 or ChR2-XXL were measured at distinct time points after injection of corresponding cRNA. Photocurrents of GtACR1 reached about 30±3% at 18 hpi, 42±10% at 42 hpi of the maximum current recorded after the normal expression time of 66 hpi. In contrast, ChR2-XXL photocurrents were 5±1% at 18 hpi, 30±8% at 42 hpi and 47±7% at 66 hpi relative to the photocurrents observed in case of GtACR1 (**Fig. S 1a**). However, the fluorescence of oocytes expressing ChR2-XXL was at least 2-times higher than of those expressing GtACR1 at all three time points though the amount of injected ChR2-XXL cRNA was four times higher (**Fig. S 1b, c**). This manifests that ChR2-XXL’s single channel conductance is strongly reduced relative to GtACR1.

As for ChRs, the functional expression of GtACR1 should depend on retinal binding ^21^. In *Xenopus laevis* oocytes, the GtACR1 photocurrent at 16 hpi was ~3 times higher in culture buffer supplemented with 1 μM all-trans-retinal than without (**Fig. 1c**), which is similar to the change in current observed for ChR2-XXL^6^. Since GtACR1::YFP fluorescence was not obvious at 16 hpi, the YFP signal at 64 hpi was investigated but no obvious difference could be seen (**Fig. 1d**), showing that the high single channel conductance produced a distinguishable photocurrent at such a low expression level that a difference in YFP signal is not detectable. Interestingly we also observed that the GtACR1 fluorescence accumulated at distinct spots (**Fig. S 2b**).

### Neurotransmission induced with ChR2-XXL

To functionally validate the powerful optogenetic tool ChR2-XXL ^6^ in primary rat cortical neurons, the DNA of ChR2-XXL was delivered into neurons using electrofection on the day of preparation. Very clear YFP fluorescence was detected on DIV13 (**Fig. 2a**) indicating ChR2-XXL expression. Fluorescence was generally homogeneous over the soma and neurites indicating an even distribution of the channel.

High photocurrents and light sensitivity make possible easy monitoring of neurotransmission, therefore we applied ChR2-XXL in primary rat cortical neurons and customised a protocol for neurotransmission studies (**Fig. 2b**). For this, neurons exhibiting fluorescence were selected as putative presynaptic neurons. Neurons without fluorescence were assumed to be postsynaptic neurons. A ChR2-XXL-expressing neuron (the putative presynaptic neuron) was photostimulated, while a ChR2-XXL negative neuron (the putative postsynaptic neuron) was recorded using whole cell patch clamp. In order to laser stimulate with high-fidelity, the distance between the putative presynaptic and postsynaptic neurons was longer than 200 μm but located in the same 0.15 mm^2^ frame of imaging. All measurements were done in the dark to avoid activation of ChR2-XXL by stray light. The principle is that neurotransmitter release from the ChR2-XXL positive presynaptic neuron under the blue light stimulus could elicit a detectable postsynaptic current (PSC) in the putative postsynaptic neuron. If excitatory, this can make the postsynaptic neuron surpass its threshold and fire action potentials.

**Figure 2.**
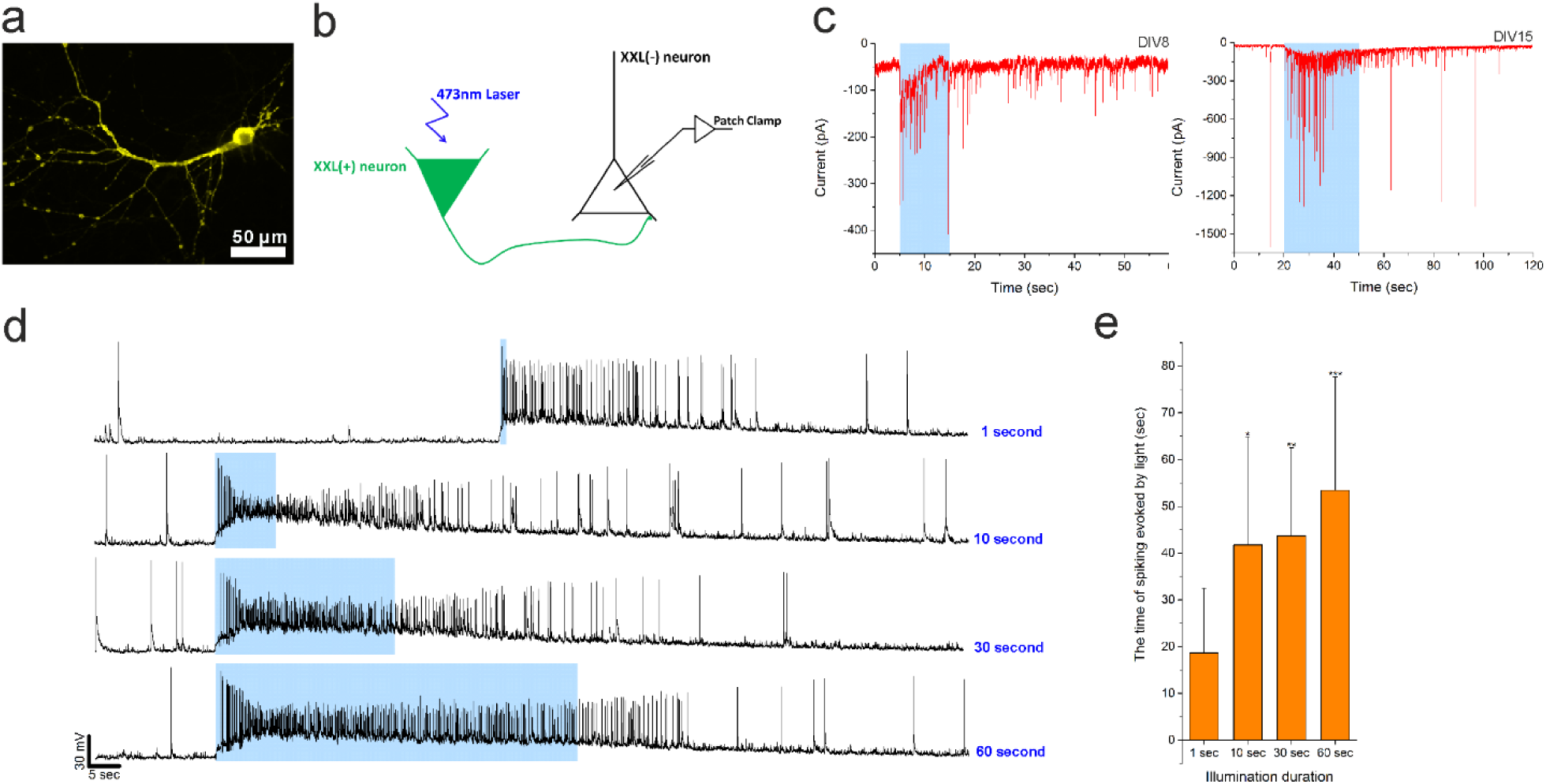
Neurotransmission induced with ChR2-XXL. a) Expression of ChR2-XXL-YFP in a primary rat cortical neuron on DIV13. b) Schematic of a neurotransmission measurement. Patch clamp was employed to measure the response of the ChR2-XXL negative neuron (the putative postsynaptic neuron) when the 473 nm laser illuminated the putative presynaptic neuron that expressed ChR2-XXL. ChR2-XXL positive neurons were identified by their fluorescence when selecting neuron pairs. c) Current traces show peak currents of ChR2-XXL negative neurons during laser stimulation at ChR2-XXL positive neurons (Holding potential: -70 mV). Blue bands indicate the period of laser stimulation. d) Spiking associated with neurotransmission from the putative presynaptic neuron under different durations of laser illumination. All traces were measured from the same neuron. Injected holding current was -27 pA (laser illumination 1 second) or -19 pA for others. e) The time of spiking evoked by the laser light of different durations (*n* = 11, 9, 15 and 8 cells for 1 sec, 10 sec, 30 sec, and 60 sec, respectively; one-way ANOVA, * p = 0.0127255, ** p = 0.00107, *** p = 0.000987 vs. 1 sec group).

Based on the above described protocol, inward currents were successfully evoked in putative postsynaptic neurons from DIV8 or DIV15 when illuminating the putative presynaptic neurons (ChR2-XXL positive) (**Fig. 2c**). Moreover, interestingly, a relatively high number of action potentials were recorded at the putative postsynaptic neurons (**Fig. 2d**), especially during long-duration photostimulation. Compared to laser stimulations of just 1 second, the postsynaptic spiking continued significantly longer when the laser stimulated for 60 seconds (**Fig. 2e**). These data indicate that neurotransmission could be shaped in primary rat cortical neurons using ChR2-XXL.

### Depolarisation block induced by ChR2-XXL

Because ChR2-XXL could generate the largest photocurrents of all ChRs published, and showed light sensitivity more than 10,000-fold over wild-type ChR2 in *Drosophila* larvae ^6^, we investigated whether ChR2-XXL could induce a stable depolarisation block when exposed to weak white light for a few minutes. In order to achieve depolarisation block, all neurons were illuminated 10 minutes with weak white light (<8.5 nW/mm^2^ from a xenon bulb). Subsequently, patch clamp recordings were made. After white light exposure, ChR2-XXL positive neurons needed more injected current than control neurons to hold the membrane potential at -70 mV regardless of DIV (**Fig. 3a**). The reason for this should be a change in the resting potential of these ChR2-XXL positive neurons caused by the continued currents through ChR2-XXL. Next, we measured the resting potential using patch clamp in current-clamp mode, with no current applied. The resting potentials of most of ChR2-XXL positive neurons were higher than those of control cells across different culture days (DIV7 to DIV14) (**Fig. 3b**). From our recordings with primary rat cortical neurons, a depolarisation to about -50 mV evoked an action potential during electrical stimulation. However, most of the ChR2-XXL positive neurons surpassed this threshold after exposure to dim white light (**Fig. 3b**). Depolarisation block drives the membrane potential over threshold and holds it there.

**Figure 3.**
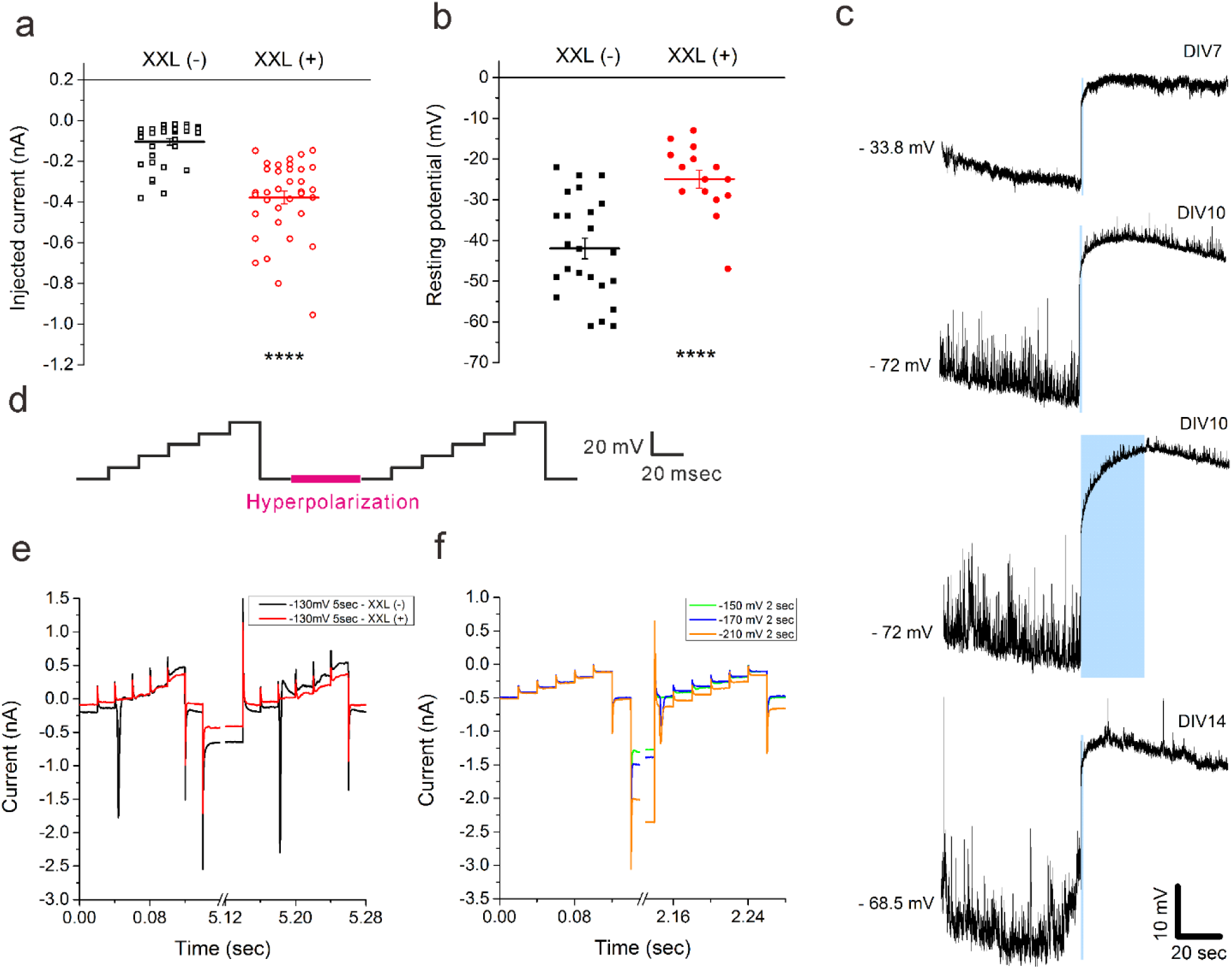
Depolarisation block induced by ChR2-XXL. a) After 10 minutes exposure to white light, ChR2-XXL positive neurons (red unfilled circles) needed more injected current than control neurons (ChR2-XXL negative neurons, black unfilled circles) to hold the membrane potential at -70 mV (DIV7 to DIV17). b) The resting potential (0 pA injection) of ChR2-XXL positive neurons (red filled circles) was higher than control cells (ChR2-XXL negative neurons, black filled circles) on DIV7 and DIV14. In both a) and b), each circle is the average value over 1 second for one neuron (t-Test, ****: a) p = 5.5233E-06, b) p = 2.72385E-10). c) No action potentials of ChR2-XXL(+) neurons are induced by laser light when directly targeted. Membrane potentials were measured with current clamp at DIV7 (laser stimulation 1 sec, injected current: -16 pA), DIV10 (laser stimulation 1 sec, injected current: -190 pA; laser stimulation 30 secs, injected current: -170 pA), DIV14 (laser stimulation 1 sec, injected current: -173 pA). d) Schematic of hyperpolarisation protocol to recover from depolarisation block. Voltage steps from -70 mV to -20 mV in 10 mV increments were applied before and after hyperpolarisation. Hyperpolarisation with different voltages and durations is indicated by the pink region e) Recovery of ChR2-XXL positive neurons was not achieved by hyperpolarisation to -130 mV for 5 secs. A typical current trace of ChR2-XXL positive neurons is shown in red, while that of a non-transfected neuron is shown in black. F) Recovery of ChR2-XXL positive neurons was successfully achieved by hyperpolarisation at the severe conditions of -170 mV for 2 secs (Blue) and -210 mV for 2 secs (Orange). These traces are from a different neurons than **Fig. 2e**.

Subsequently we investigated whether the membrane potential could be depolarised in ChR2-XXL positive neurons by laser stimulation after a depolarisation block. Together with a laser stimulus, patch clamp was employed to characterise ChR2-XXL positive cortical neurons. All representative voltage traces indicated two kinds of patterns (**Fig. 3c**). The first pattern showed a declining membrane potential prior to laser stimulation due to exposure to the room lighting while setting up the experiment, which demonstrated ChR2-XXL had been activated but was gradually closing. This is consistent with ChR2-XXL’s long open state and high light sensitivity ^6^. The second pattern was that depolarisation took place without action potentials during laser stimulation of ChR2-XXL-expressing neurons, indicating that the voltage-gated channels were still inactivated. Moreover, prolonged laser stimulation failed to elicit action potentials. These data demonstrated that ChR2-XXL can be used to induce a stable state of depolarisation block.

A depolarisation block forces neurons into a refractory state without the ability to produce action potentials due to inactive voltage-gated sodium channels ^13^. Therefore, we designed an approach using voltage clamp to measure how voltage-gated inward current was changed and confirmed that it could be recovered in ChR2-XXL positive neurons after neurons were blocked. To recover blocked neurons (**Fig. 3d**), voltage was stepped from -70 mV to -20 mV by +10 mV steps each held for 20 msec. Then a hyperpolarisation pulse was applied and those steps repeated. Different hyperpolarisations were tested as in Bendahhou ^22^. No inward current was detected before or after hyperpolarisation at -130 mV for 5 secs in ChR2-XXL positive neurons, whereas the amplitude of voltage-gated inward current was slightly increased after hyperpolarisation in a control neuron (**Fig. 3e**). These data indicated that hyperpolarisation can un-block a control neuron, but those particular conditions were not sufficient for un-blocking the ChR2-XXL expressing neuron. Therefore we further decreased the hyperpolarizing pulse for ChR2-XXL positive neurons. Finally, at -170 mV for 2 secs or -210 mV for 2 secs, a current recovery was seen, whereas it never appeared at -150 mV for 2 secs (**Fig. 3f**). Moreover, compared with the hyperpolarisation at -170 mV for 2 secs, voltage-gated current was slightly increased after hyperpolarisation at -210 mV, but it was still lower than the voltage-gated current in the control neurons. These data indicated that depolarisation block could be successfully induced using ChR2-XXL and that it could be recovered by hyperpolarisation to -170 mV or -210 mV for 2 secs.

### Light-gated chloride conduction by GtACR1 during neural development

Having successfully induced neurotransmission and a depolarisation block in primary rat cortical neurons using a depolarizing tool, we aimed to apply the high current, light-gated, anion channel GtACR1 to silence neuronal activity by hyperpolarisation. Light-gated proton pumps ^23^ or modified ChRs with additional chloride conductance ^24–27^ are rather inefficient compared to optogenetic channels. GtACR1-YFP DNA was transfected into primary rat cortical neurons by electrofection before seeding cells on PDL-coated coverslips. Very clear yellow fluorescence was detected in the neurons transfected with GtACR1 on DIV16 (**Fig. 4a**). However, fluorescence was intermittent along the neural membrane including the membrane of axons or dendrites. These data indicated GtACR1 expression in the membrane of neurons.

**Figure 4.**
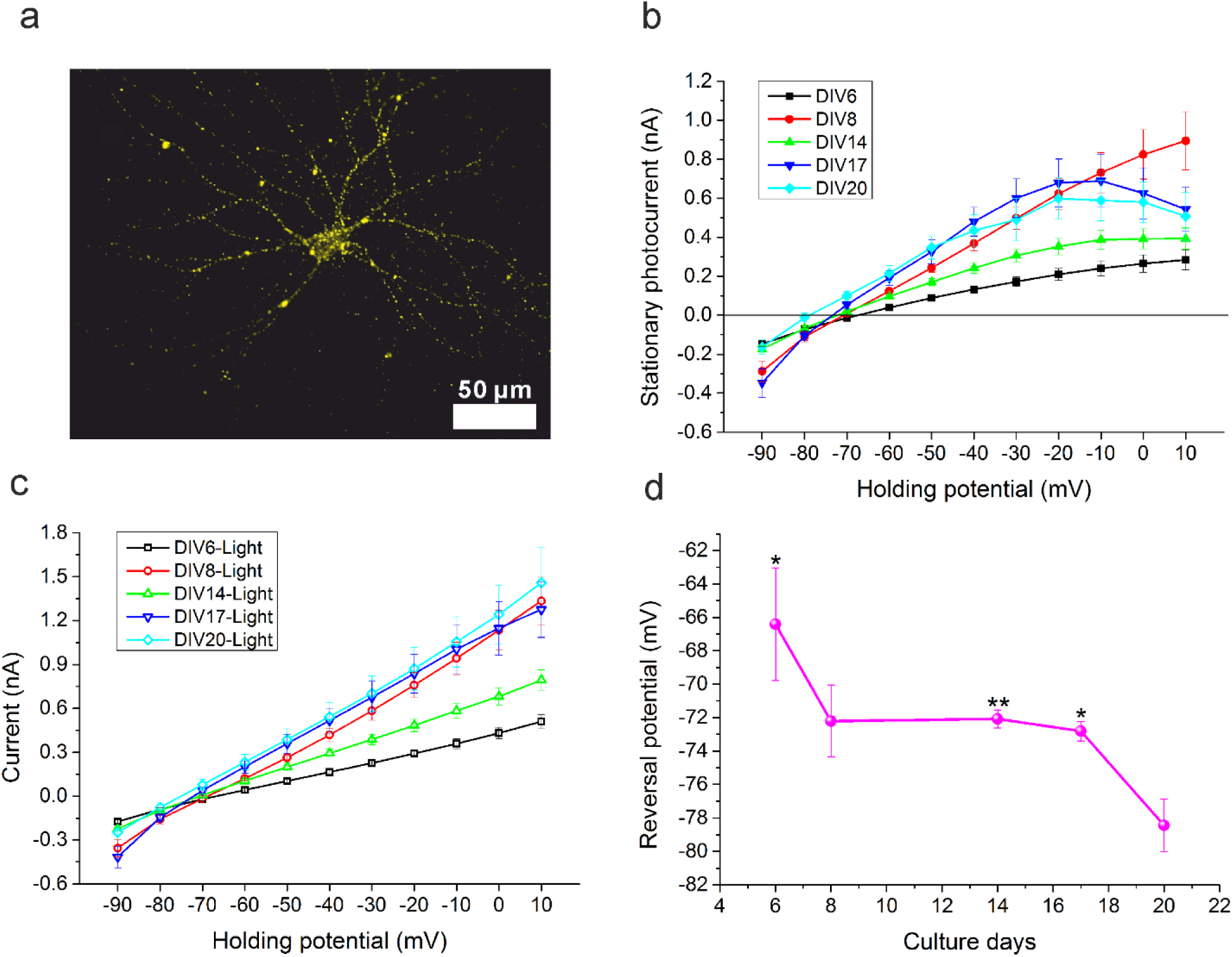
IE relationship for GtACR1 in primary rat cortical neurons. a) Yellow fluorescence is shown in a typical rat cortical neuron (DIV16) transfected with GtACR1-YFP. b) IE relationship for GtACR1 in primary rat cortical neurons at different days. The data (mean values ± s.e.m.; *n*= 3 to 5 neurons) were baseline corrected by subtracting the current from the last 50 milliseconds without light from the current of the last 50 milliseconds with light. c) The current of GtACR1-expressing neurons clamped at different voltages from -90 mV to 10 mV and exposed to blue light. The data indicates mean values of the last 50 milliseconds ± s.e.m. (*n*= 3 to 5 neurons). d) Reversal potential for GtACR1 during neural development. The reversal potential was defined as the point where the stationary photocurrent amplitude was 0 pA. At different days in *vitro*, the reversal potentials of GtACR1 (Mean values ± s.e.m.; *n* =3 to 5 cells) were determined by linear fits of the IE relationship curves as depicted in **Fig. 4b**. Data were analysed by one-way ANOVA (* p = 0.017878 DIV6 vs. DIV20 group, p = 0.032382 DIV17 vs. DIV20 group, ** p = 0.00858 DIV14 vs. DIV20 group).

Next, the function of GtACR1 was investigated by electrophysiology and light induced opening. GtACR1 can be activated with a saturating blue light pulse (**Fig. S 2a**), so here blue laser was employed to investigate GtACR1 in primary cortical neurons. The membrane of cortical neurons was hyperpolarised by blue laser stimulation for 0.25 milliseconds (**Fig. S 4a**), hyperpolarisation was induced by GtACR1 with a mean rise time (π_R_) of 1.23 ± 0.14 milliseconds and a mean decay time (π□) of 126.03 ± 19.43 milliseconds (**Fig. S 4b**). This is similar to slow switching components observed in GtACR1 by Sineschchekov et al [33]. Voltage step sequences applied during and 1.5 s after illumination exhibited lower currents at each holding potential. Moreover, voltage-gated peak current was eliminated by laser stimulation (see: inset in **Fig. S 5a**). At early stages of neuronal development (DIV6 and DIV8), the IE relationship showed a linear relationship (**Fig. 4b**). However, at later, more mature states of neurons (DIV14, DIV17 and DIV20) a rectification appeared. Therefore, we investigated whether the synaptic inputs in the mature neural network resulted in this rectification. The later mature state of neurons (DIV14, DIV16 and DIV20) still showed similar rectification behaviour on stationary photocurrents (**Fig. S 6**). Therefore, the rectification was not associated with synaptic inputs in the neuronal network. Next, we compared the voltage-dependent currents during photostimulation and dark-periods. Currents during photostimulation fit to a linear shape (**Fig. 4c** and **Fig. S 5 c**), whereas these currents under dark showed a rectification behaviour (**Fig. S 5b** and **d**), which contributed to the age-dependent rectification behaviour of the stationary photocurrent (**Fig. 4b**). In a word, the change in rectification is not associated with the GtACR1 channels, but the state of the neurons in which they are expressed.

Furthermore, the reversal potential for GtACR1 during neural development (from DIV6 to DIV20) showed two kinds of trends (**Fig. 4d**). For younger neurons from DIV6 to DIV8, the reversal potential declined from -66.4 mV ± 3.4 mV to -72.2 mV ± 2.1 mV. The reversal potential was then steady through DIV17 at about -72 mV (p = 0.958 for comparisons of DIV8 to DIV14 and p = 0.8417 for DIV8 to DIV17. Then at DIV20, it further declined to -78.4 mV ± 1.6 mV (p = 0.032, DIV17 vs. DIV20). These data imply the chloride concentration in neurons was changing during neural development.

### Photoinhibition of spiking by GtACR1

Having successfully validated the light-gated chloride conductivity of GtACR1 in primary rat cortical neurons, we investigated how GtACR1 mediated neuronal membrane potential alterations. First, we addressed photoinhibition of spiking induced by pulsed currents. To induce spiking, 10 millisecond long 300 pA depolarizing electric pulses were applied at 20 Hz to primary rat cortical neurons expressing GtACR1 via the patch clamp pipette in whole cell mode. For a typical GtACR1-expressing neuron on DIV8, illumination successfully inhibited induced neuronal spiking compared to current injections performed in the dark (black trace vs. green trace, see: **Fig. 5a**). Also, the same results were obtained at DIV14 (**Fig. 5b**) when neurons are embedded in a neural network and should also be receiving synaptic inputs. Because the chloride concentration in neurons was different during neural development (**Fig. 4d**), we wanted to verify whether GtACR1 could inhibit neural activity at different culture days. Therefore we tested 4-7 neurons at each different culture day (from DIV6 to DIV20). Interestingly, all spikings induced by pulsed currrent were successfully inhibited by light, regardless of the neuronal age and correspoding chloride concentrations. These data indicate that the large currents generated by GtACR1 overcome neural stimulation even when chloride concentration reduces flow through the channel.

**Figure 5.**
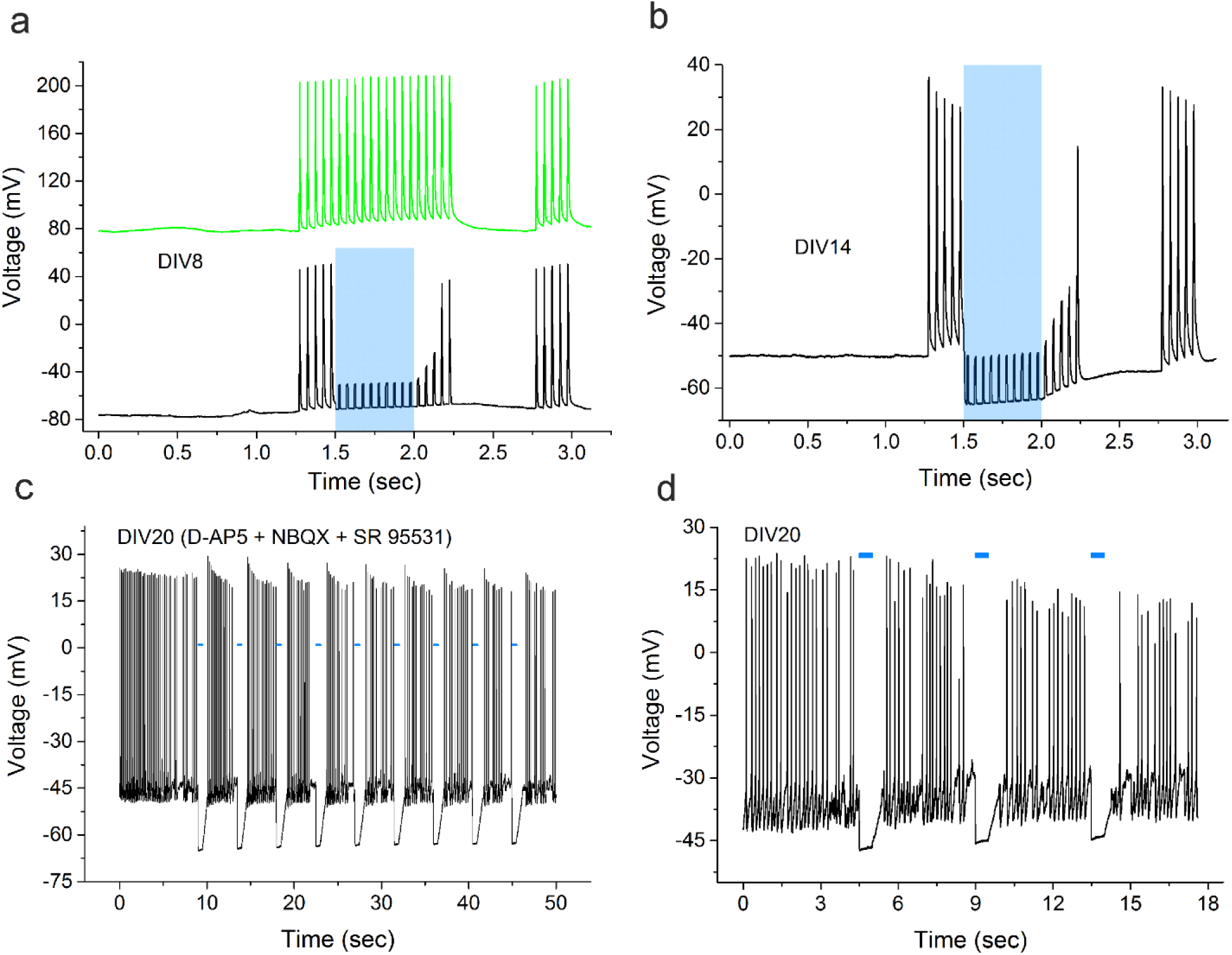
Photoinhibition of primary rat cortical neural activity at different culture days. a) Photoinhibition of spiking induced by pulsed current (300 pA for 10 milliseconds in one pulse, 20 Hz) in a typical GtACR1-expressing neuron of DIV8. Green trace indicates the meausurement without light (top), black trace (bottom) indicates the measurement with light for 500 msec (blue box). b) Photoinhibition of spiking induced by pulsed current (300 pA for 10 milliseconds in one pulse, 20 Hz) in a typical GtACR1-expressing neuron at DIV14. c) The amplitude of action potentials increased after photoinhibition. In a typical neuron (DIV20) expressing GtACR1, a continuous current of 100 pA was clamped near threshold so it continually fired APs. Synaptic blockers were used in the bath. Nine blue laser pulses were applied for inhibiting neural activity, each pulse lasted for 500 milliseconds separated by a delay time of 4 seconds and showed higher amplitude APs when the firing resumed in the dark. d) Photoinhibition of spontaneous activity in a GtACR1-expressing neuron at DIV20 without any current injection or synaptic blockers. Each pulse of blue laser lasts 500 msec (blue bar). In all measurements, blue laser intensity was 0.21 W/mm^2^.

Subsequently, we further addressed spontaneous neural activity in mature networks. Neurons expressing GtACR1 at DIV 20 were selected for whole cell patch clamp recordings. In response to photostimulation, spontaneous activity of all neurons measured (*n*=10 neurons expressing GtACR1 from DIV20) was successfully inhibited (**Fig. S 7**). Furthermore, spontaneous activity was repeatly suppressed with intervening return to spontaneous firing in the dark (**Fig. 5d**). These data demonstrate GtACR1 could effectively and reversibly inhibit spontaneous neural activity by illumination.

Furthermore, whether hyperpolarisation induced by GtACR1 affects the spiking after illumination is still unknown. In order to address this open gap, during the recordings of GtACR1-expressing neurons a continuous depolarizing injection of 100 pA was applied to generate continuous action potentials. Simultaneously, laser pulses were applied to induce hyperpolarisation. The amplitude of action potentials after light illumination was slightly increased (**Fig. 5c**). Demonstrating that the hyperpolarisation by GtACR1 could recover the voltage-gated channels in neurons, resulting in increasing the action potential’s amplitude (**Fig. 5c**).

However, by multiple illuminations, the frequency of action potentials showed a decreasing phenomenon. This may be due to after turning off the light inhibition the cells rebound and fire too much, and too much firing triggers more slow calcium activated outward K^+^ currents as a response to over-excitation ^14,28^. Also, we tested whether the amplitude of hyperpolarisation by GtACR1 is changed after multiple activations. The photocurrent evoked by the laser pulse decreased to 75% after ~ 10 pulses (**Fig. S 8a and b**). Neural membrane potential was hyperpolarised by multiple light pulses (**Fig. S 8 c and d**) and interestingly, the hyperpolarisation amplitude decreased following multiple pulses due to GtACR1’s desensitisation property.

## Discussion

The high photocurrents of GtACR1 and ChR2-XXL that have been shown here and previously in X.l. oocytes is attributable to different mechanisms. High currents reached for GtACR1 with low cRNA injection and expression levels below the fluorescence detection limit suggest higher single channel conductance. Whereas, ChR2-XXL reached high currents by producing twice as many channels, according to fluorescent tag intensity. The rapid establishment of detectable GtACR1 currents in X.l. oocytes provides a platform for the further study of GtACR1 by site-directed mutagenesis. The aim of these efforts will be to generate mutants with faster kinetics and a red-shifted absorbtion spectrum.

ChR2-XXL activation was used to control non-expressing putative postsynaptic cells. Interestingly, the spiking of putative postsynaptic neurons persisted after the end of a stimulus (**Fig. 2e**) with a slow decline. This phenomenon reflected the long open-state of ChR2-XXL. In addition, the amplitude of action potentials declined during the illumination (**Fig. 2d**). After laser illumination and a short dark period, the amplitude of action potentials increased to the initial size observed during spontaneous activity. This phenomenon implies that synaptic inputs elicited by the large and sustained photocurrent of ChR2-XXL, had blocked the recovery of some sodium channels from inactivation, resulting in reduced action potentials. Furthermore, the spiking in these postsynaptic neurons showed an oscillating decline (**Fig. S 3**). We hypothesise that the observed oscillating decline is associated with ChR2-XXL inducing rhythmic patterns of calcium influx in the presynaptic neurons, which can trigger exocytosis of neurotransmitter-containing synaptic vesicles via mechanisms shown previously ^29^. The photocurrent of ChR2-XXL shows a prolonged decay ^6^, which is presumably responsible for the decline of the spike rate.

With sustained current injection, neurons eventually enter a depolarisation block, which can protect them from excessive spiking activity ^30^. Here, we have shown that ChR2-XXL can be used to induce a depolarisation block (**Fig. 3c**) by exposing the neurons to a halogen lamp’s white light for 10 minutes. After this, all voltage traces (**Fig. 3c**) showed a decreased potential in advance of the first laser photostimulation, indicating that ChR2-XXL channels were still not completely closed. White light with low intensity is able to keep ChR2-XXL channels open, resulting sustained depolarization. After 10 minutes exposure to white light, our results showed that photostimulation failed to provoke action potentials (see: **Fig. 3c**). These data suggest that because the peak current and sensitivity are so high, even using the tail of the absorption spectrum, it may be possible to manipulate ChR2-XXL with longer wavelength light that is more favoured for *in vivo* applications. Moreover, it was difficult to recover this state by hyperpolarisation at lower holding voltages. The need for at least -170 mV hyperpolarisation to re-set voltage gated channels suggests most natural processes would fail to overcome manipulation by ChR2-XXL. Our results support ChR2-XXL as a potential tool to block highly excitable neurons, as seen in some disorders like peripheral neuropathic pain.

To block neural firing by the more classical approach of hyperpolarization GtACR1 was expressed in primary cortical neurons. Contrary to the expectation that older neurons would have produced more protein and therefore have higher photoinduced currents, the response at DIV8 was similar to DIV17 or DIV20 (**Fig. 4c**). It may be that the CMV promoter is only stable in neurons of particular types ^33^, independent of the transgene, since during neuronal development GtACR1 shows normalised currents with light that are not significantly different (**Fig. S 5c)**. The recombinant Adeno-Associated Virus 6 (rAAV6) with hSyn promoter system would eliminate this issue in primary rat cortical neurons by utilizing a continually active neuron-specific promoter. The use of rAAVs would furthermore increase the percent of neurons expressing GtACR1 ^34^. Another factor influencing the current over time may be the ECl changes that have been shown to occur in the developing rodent brain ^35^. Figure 4d implies chloride concentration was decreasing during neural development, consistent with other studies ^35,36^. It may therefore be possible to use GtACR1 to manipulate chloride concentration during cell maturation to further study these processes.

## Methods

### Molecular biology

The ChR2-XXL-YFP fragment was amplified through Polymerase Chain Reaction with Phusion High-Fidelity DNA Polymerase (2 U/µL, # F-530S, Thermo Scientific, Germany) based on the plasmid named pGEM ChR2-XXL-YFP. Primer pairs for its amplification are listed below (**Table 1**). For ChR2-XXL subcloning, the psc-hSyn-ChR2-XXL was generated using restriction digest-based strategies ^37^, with the backbone produced from psc-hSyn-ChR2opt ^17^ using double-digest with *Hind* III and *Bam* HI.

**Table 1.**
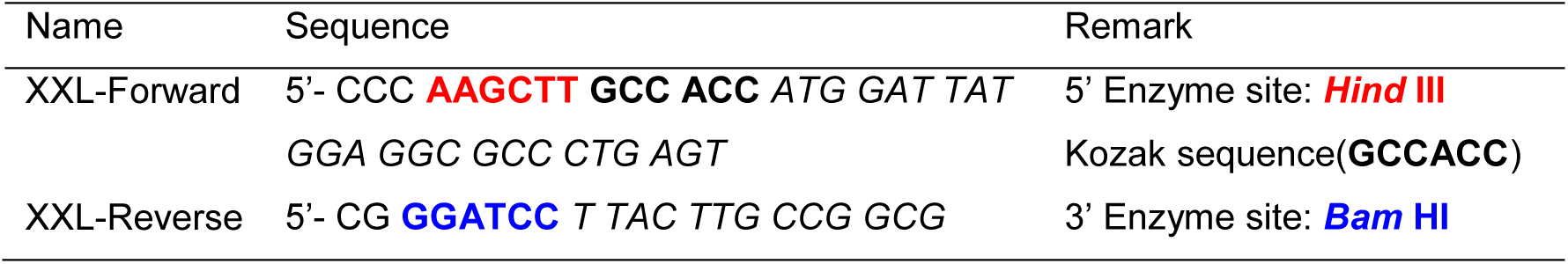
Primer pair for amplification of ChR2-XXL-YFP

5’ Enzyme site is marked in red colour, 3’ Enzyme site is marked in blue colour. Kozak sequence ^38^ improving efficient initiation of translation is marked in black colour with bold font.

The GtACR1 sequence was synthesised by GeneArt Strings DNA Fragments (Life technologies, Thermo Fisher Scientific) according to the publication ^7^, while the codon usage was optimised to *Drosophila Melanogaster*. For *Xenopus* oocyte expression, the GtACR1 DNA was inserted into the pGEMHE vector within N-terminal BamHI and C-terminal XhoI restriction sites; a YFP tag was fused after the XhoI restriction site. For cortical neuron expression, the GtACR1-YFP sequence was cut from pGEMHE-GtACR-YFP and inserted into the PBK CMV vector using N terminal BamHI and C terminal HindIII restriction sites. The constructs were then confirmed by sequencing (GATC, Konstanz).

### Maxi preparation of Plasmids

A single colony was picked from a freshly streaked selective plate and used to inoculate a starter culture of 5 ml LB medium containing the antibiotic Ampicillin (50 μg/mL). The starter culture was incubated for approx. 8 h at 37°C with vigorous shaking (approx. 225 rpm). 200 ml (pBK-CMV-GtACR1-YFP) or 500 ml (psc-hSyn-ChR2-XXL due to its low-copy) medium were inoculated with 250-500 μl of starter culture. Then cultures were grown at 37°C for 12-16 h with vigorous shaking (approx. 225 rpm). To purify plasmids for electroporation, EndoFree Plasmid Maxi Kit (#12362 Qiagen, Germany) was used following the protocol the kit provided. In the last step, the DNA was dissolved in 200 μL of TE buffer. To determine the yield, DNA concentration was determined by both the microvolume UV-Vis spectrophotometry (NanoDrop 1000, Thermo Fisher Scientific, MA, USA) at 260 nm and quantitative analysis on an agarose gel after electrophoresis.

### Primary Rat Cortical Neuron culture and Electrofection

Primary cortical cultures were prepared as described previously ^39^. Briefly, cortices from embryonic day 18 (E18) Wistar rat brains were dissected and mechanically dissociated by trituration with a fire-polished, silanized pasteur pipette in 1 ml Hank’s balanced salt solution without calcium or magnesium (HBSS-) (0.035% sodium bicarbonate, 1 mM pyruvate, 10 mM HEPES, 20 mM glucose, pH 7.4). The cell suspension was diluted 1:2 in HBSS+ (with calcium and magnesium) and non-dispersed tissue was allowed to settle for 3 min. The supernatant was centrifuged for 2 min at 200 g. Amaxa Rat Neuron Nucleofector^®^ Kit (Cat.No. VPG-1003) from Lonza Cologne GmbH (Germany) was used to transfer circular, double stranded, plasmid DNA directly into cells. Each electrofection experiment used two fresh cortices, dissociated and collected as described above. After centrifugation for 2 min at 200 g, the cell pellet was carefully resuspended in 200 µl room temperature Nucleofector^®^ Solution from the Kit (163.6 µl of Rat Neuron Nucleofector^®^ Solution mixed with 36.4 µl of Supplement), to which was added 6 µg of psc-hSyn-ChR2-XXL plasmid or 8.5 µg pBK-CMV-GtACR1-TYE plasmid. The cell/DNA suspension was quickly transferred into a certified cuvette without producing bubbles. Using the Nucleofector^®^ Device (AAD-1001S, Lonza, Germany), program G-013 was applied to the cells. 800 μl of the pre-equilibrated RPMI medium (Gibco, Grand Island, NY, USA) with 1% FBS (Gibco, Grand Island, NY, USA) and 0.5 mM L-glutamine (Gibco, Paisley, UK) were immediately added to the cuvette The cells were counted in a Neubauer counting chamber and plated in a concentration of 50,000 cells per well (24-well-plate) on poly-D-lysine (PDL, 10 µg/ml) coated, 12 mm diameter coverslips in 500 μl supplemented RPMI medium per well. After 4 hours, the medium was carefully replaced with 500 μl fresh supplemented Neurobasal medium (NB) per well to remove cellular debris (supplemented Neurobasal medium (Gibco, Paisley, UK) with 1% B-27 (Gibco, Grand Island, NY, USA), 0.5 mM L-glutamine (Gibco, Paisley, UK) and 50 µg/ml gentamicin (Sigma, Steinheim, Germany)). Cells were kept at 37°C, 5% CO_2_ and 100% humidity in 500 µl NB. Every third day, half of the medium was exchanged.

This work except for GtACR1 experiments was carried out with the approval of the Landesumweltamt für Natur, Umwelt und Verbraucherschutz Nordrhein-Westfalen, Recklinghausen, Germany, in accordance with §6 TierschG., §4 TSchG i.V. and §2 TierSchVerV. GtACR1 experiments were carried out with the approval of the Landesumweltamt für Natur, Umwelt und Verbraucherschutz Nordrhein-Westfalen, Recklinghausen, Germany, number 84-02.04.2015.A173.

### Fluorescence Microscopy and quantification of homogenised oocytes

To qualitatively monitor the expression of YFP-tagged ionotropic photoreceptors conventional fluorescence microscopy was used. The YFP fluorescence of oocytes was detected by Leica DMi8. Bright field settings for the inverted microscope were always 100 ms exposure and gain 1.5. Corresponding settings for detection of YFP fluorescence were always 1600 ms exposure and gain 1.8.

To quantify the fluorescence intensity, *Xenovus* oocytes expressing the indicated constructs were homogenised by pipetting and the resulting lysate was loaded into Nunc surface 96-well plates and investigated using a corresponding fluorometer. The fluorescence emission was then measured at 538 nm by a Fluoroskan Ascent microplate fluorometer with 485 nm excitation.

### Oocyte electrophysiology

cRNA was generated with the AmpliCap-MaxT7 High Yield Message Maker Kit (Epicentre Biotechnologies) using NheI-linearised pGEM-HE GtACR1 YFP plasmid. Oocytes were injected with the indicated amount of cRNA and incubated in ND96 solution (96 mM NaCl, 2 mM KCl, 1 mM MgCl_2_, 1 mM CaCl_2_, 10 mM HEPES and 50 ng/ml Gentamycin, pH 7.4) containing 1 μM ATR. Ringer’s solution (110 mM NaCl, 5mM KCl, 2 mM BaCl_2_, 1 mM MgCl_2_, 5 mM HEPES, pH 7.6) was used as standard buffer for Two electrode voltage-clamp recordings with the indicated holding potential. 2s 473 nm blue laser and 530 nm green laser saturating light pulses were used for illumination.

### Neuron electrophysiology

Whole cell patch clamp experiments were performed using an EPC9 amplifier (HEKA Elektronik, Lambrecht/Pfalz, Germany) controlled by the TIDA 5.05 software (HEKA Elektronik). The recordings were performed in extracellular patch clamp buffer solution (mM: NaCl 120, KCl 3, MgCl_2_ 1, HEPES 10, CaCl_2_ 2; pH 7.3). For the synaptic input blocking measurements, the following (all from Tocris Bioscience, Bristol, UK) were added to the extracellular solution above: D-AP5 (25 µM), NBQX (10 µM), SR 95531 hydrobromide (alternative name: Gabazine, 20 µM). Patch pipettes were pulled from borosilicate glass capillaries (Sutter Instrument Co., Novato, CA, USA) using a micropipette puller (P-2000, Sutter) to achieve a pipette resistance of 6-9 MOhm. For ChR2-XXL measurements the pipettes were filled with intracellular patch solution (mM: NaCl 2, KCl 120, MgCl_2_ 4, HEPES 5, EGTA 0.2, Mg-ATP 0.20, pH 7.3). For GtACR1 measurements, an intracellular solution with low chloride concentration (mM: K_2_SO_4_ 67.5, MgCl_2_ 2, adjusted to 248 mosm/kg with glucose) was used.

### Laser stimulation

Optogenetic channel expressing neurons were stimulated with a 473 nm diode laser (Rapp OptoElectronic GmbH, Hamburg, Germany), which was guided through an optical fibre and reflected with a dichroic mirror (UGA-40, Rapp OptoElectronic GmbH) into the light path of a Zeiss Axioscope microscope equipped with a x20 water immersion objective (Olympus, UMPLFLN20xW). Different delay times (500 msec to 60 sec) and different laser pulse times (ranging from 0.25 msec to 100 sec) were applied for light stimulation. Light intensity was 1.68 W/mm^2^ for ChR2-XXL stimulation (laser spot diameter was about 24 µm); GtACR1 can also be activated with a saturating blue light pulse (**Fig. S 2a**), light intensity was 0.21 W/mm^2^ for GtACR1 (laser spot diameter was about 61.2 µm).

### Data analysis

Data were analysed in TIDA 5.240, Excel, Matlab, or Origin 9. The data are given as mean ± standard error of the mean (neurons) or s.d. (oocytes). For ChR2-XXL data, the spikes whose amplitudes were more than 35 mV were selected to determine the oscillation of firing frequency. Each frequency point was the average value of 5 seconds window. For GtACR1 measurements, stationary photocurrents of GtACR1 were measured for the last 50 ms of the illumination period, and then were corrected by subtraction of the current values from the last 50 milliseconds without light to adjust the baseline to zero. Reversal potentials were calculated by linear fit between a measured positive and a measured negative stationary current to extract the I=0 pA crossing point (here between two data points at voltage step of -60 mV or -80 mV and -70 mV).

## Data Availability

The datasets generated during and/or analysed during the current study are available from the corresponding author on reasonable request.

## Acknowledgements

The authors would like to thank B. Breuer for preparation of primary neurons.

## Author Contributions

Conceptualisation: LJ, SG, AO. Data collection: LJ, EFJ, SG, WL. Data curation: SG, AO. Formal analysis: LJ, EFJ, SG, VM. Supervision: SG, VM, AO. Visualisation: LJ, EFJ, VM, SG, AO. All authors have contributed to and read the final version of the manuscript.

